# Genetic variance in the murine defensin locus modulates glucose homeostasis

**DOI:** 10.1101/2024.07.25.605202

**Authors:** Stewart W.C. Masson, Rebecca C. Simpson, Harry B. Cutler, Patrick W. Carlos, Oana C. Marian, Meg Potter, Søren Madsen, Kristen C. Cooke, Niamh R. Craw, Oliver K. Fuller, Dylan J. Harney, Mark Larance, Gregory J. Cooney, Grant Morahan, Erin R. Shanahan, Christopher Hodgkins, Richard J. Payne, Jacqueline Stöckli, David E. James

## Abstract

Insulin resistance is a heritable risk factor for many chronic diseases; however, the genetic drivers remain elusive. In seeking these, we performed genetic mapping of insulin sensitivity in 670 chow-fed Diversity Outbred in Australia (DOz) mice and identified a genome-wide significant locus (QTL) on chromosome 8 encompassing 17 defensin genes. By taking a systems genetics approach, we ultimately identified alpha-defensin 26 (Defa26) as the causal gene in this region. To validate these findings, we synthesized Defa26 and performed diet supplementation experiments in two mouse strains with distinct endogenous Defa26 expression levels. In the strain with relatively lower endogenous expression (C57BL/6J) supplementation improved insulin sensitivity and reduced gut permeability, while in the strain with higher endogenous expression (A/J) it caused hypoinsulinemia, glucose intolerance and muscle wasting. Based on gut microbiome and plasma bile acid profiling this appeared to be the result of disrupted microbial bile acid metabolism. These data illustrate the danger of single strain over-reliance and provide the first evidence of a link between host-genetics and insulin sensitivity which is mediated by the microbiome.

## Introduction

Insulin is among the most potent hormones in the human body; therefore, it is not surprising that defects in insulin action, such as insulin resistance (IR) are shared risk factors for many chronic diseases (1). Studies in twins and first-degree relatives of individuals with Type 2 Diabetes have provided strong evidence of a genetic component (2, 3). However, with some notable exceptions (4–6), genome-wide association studies have failed to identify loci for IR. One potential explanation is that the diversity of human environments confounds genetic signals via complex gene-by-environment interactions. One manifestation of the environment that has been implicated in metabolic disease is the gut microbiome (7–9). Intriguingly, some microbial compositions have been shown to have beneficial effects on metabolic health while others appear harmful (10). These effects are thought to be conveyed via diverse signalling molecules including peptides (11, 12), metabolites (13, 14) and bile acids (15–18). However, the host-microbiome relationship is bi-directional, as host genetics can also influence gut microbial composition. Compelling evidence of this has been provided by human genome-wide association and twin studies (19, 20). Furthermore, a recent study in mice revealed that one quarter of all detected microbial taxa have significant quantitative trait loci (QTL) implying host regulation of their abundance (21). Notably, this includes the metabolically beneficial *Akkermansia muciniphila*. However, it is unknown if genetic drivers of microbe abundance can modulate insulin sensitivity.

One mechanism for host regulation of the gut microbiota is the secretion of anti-microbial peptides called defensins into the intestinal lumen. Defensins are an ancient component of the immune system, found across the tree-of-life from plants to humans. They are small peptides (29-40 amino acids) with potent antibacterial and antiviral activities (22). In humans, α-defensins are produced by neutrophils and specialised gut epithelial cells, called Paneth cells. However, in mice, Paneth cells are the only source of α-defensins (also called cryptidins) (23). Defensin secretion is regulated in response to bacteria, nutrients or cholinergic agonists (24). Genetic variation in α-defensin expression among different mouse strains has been reported (25) although this has not been systematically examined. Defensins have also previously been linked to metabolic health. Oral administration of human α-defensin-5 to mice ameliorated liver fibrosis, diet-induced non-alcoholic liver steatosis, dyslipidaemia and glucose intolerance (26–29). There has been some interest in defensins as a therapeutic due to their oral bioavailability, a characteristic lacking in most peptide-based therapies.

Mice are a valuable tool for studying human diseases due to their genetic and physiologic similarities. Moreover, they afford the precise environmental control required to detect genetic loci associated with complex diseases. This has been illustrated by studies in panels of inbred mouse strains fed different diets (30–34) and genetically diverse mouse populations such as Jackson Laboratory’s Diversity Outbred (DO) mice (35–38). We have established a similar population of mice, which we term Diversity Outbred in Australia (DOz), and have previously used this population to investigate skeletal muscle insulin resistance (39), and the metabolic consequences of weight-cycling (40). Here, we use these mice to explore genetic drivers of insulin sensitivity. Using the Matsuda Index as a surrogate measure of whole-body insulin sensitivity, we identify a QTL within the defensin locus. Sequence variation at this locus were associated with increased expression of α-defensin 26 (Defa26) and the abundance of metabolically beneficial microbes. We validated this observation via dietary supplementation with synthetic Defa26 in C57BL/6J mice, before going on to demonstrate maladaptive effects in another strain, potentially via changes in microbial derived bile acids. These data provide insight into the genetic architecture of insulin sensitivity and highlight the limitations of testing therapeutics in single mouse strains.

## Results

### Whole body insulin sensitivity is genetically linked to the defensin locus on chromosome 8

To quantify whole body insulin sensitivity, we conducted glucose tolerance tests on 670 chow-fed male DOz mice (Figure 1A) and used these data to calculate the Matsuda Index, a surrogate measure of insulin sensitivity (41). Consistent with our previous work (39), we observed profound variation in the Matsuda Index despite all animals being housed under identical conditions and fed the same chow diet. We next performed genetic mapping of the Matsuda Index (Figure 1B) and identified 2 genome-wide significant quantitative trait loci (QTL), one on chromosome 2 at 162.2 Mbp and a second on chromosome 8 at 21.7 MBp. The chromosome 8 locus centred over the defensin gene cluster, a syntenic region shared between mice and humans (42), which contains 53 defensin genes and 22 defensin pseudogenes (Figure 1C). Considering previous links between defensins and metabolic disease (26–29) we selected this QTL for further validation.

**Figure 1.**
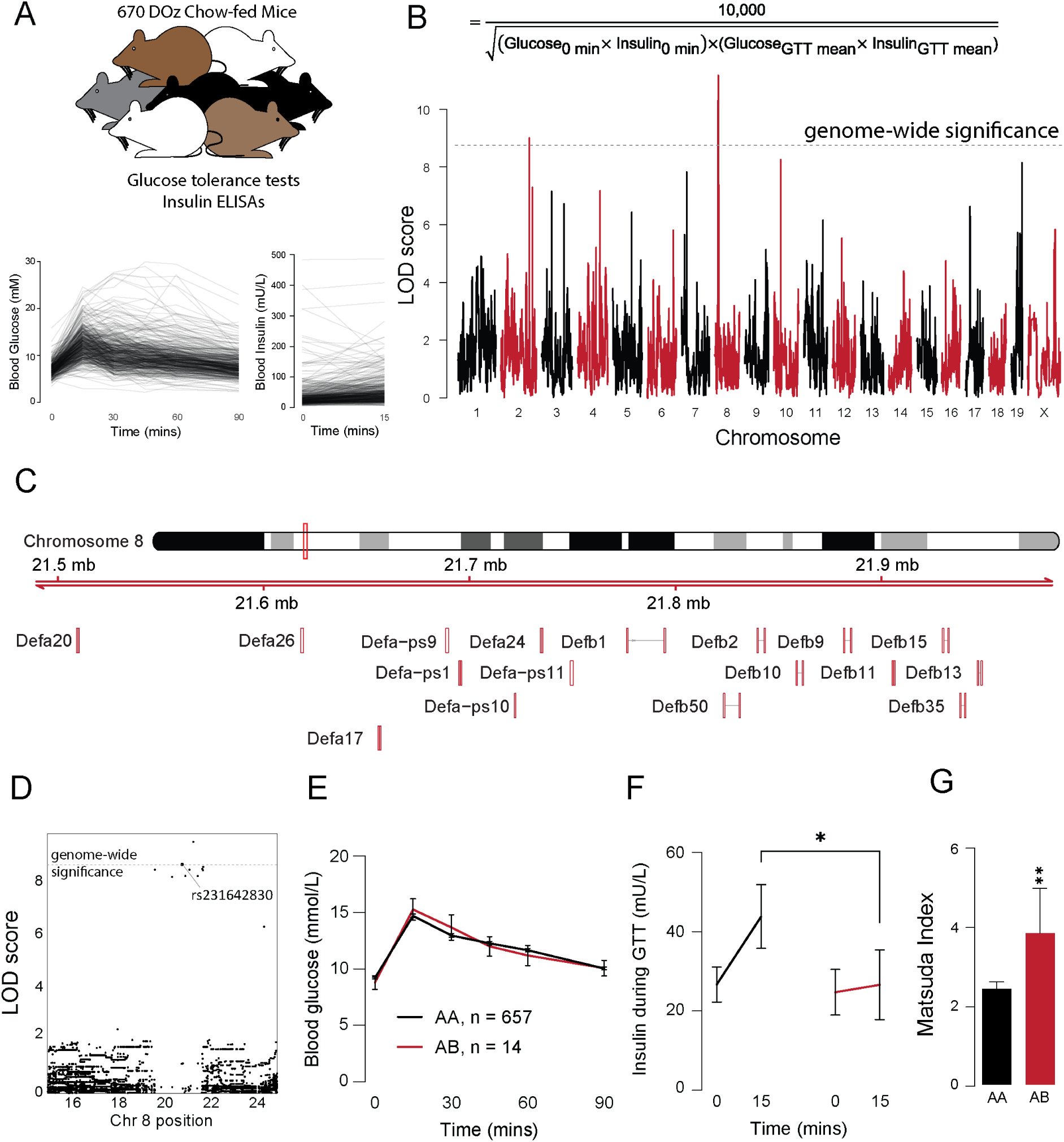
– Genetic mapping of insulin sensitivity in 670 chow-fed Diversity Outbred in Australia mice. **A)** Glucose and insulin concentrations during a glucose tolerance test in 670 DOz mice. **B)** The equation used to calculate the Matsuda Index and genetic mapping of the Matsuda Index in DOz mice. **C)** The defensin locus on chromosome 8. **D)** Single nucleotide polymorphism (SNP) mapping of the Matsuda Index on chromosome 8. **E)** Blood glucose concentrations during a glucose tolerance test of mice carrying the minor allele of rs231642830 (AB) and non-carrier (AA) controls. **F)** Blood insulin concentrations during a glucose tolerance test of mice carrying the minor allele of rs231642830 (AB) and non-carrier (AA) controls. **G)** Matsuda Index of mice carrying the minor allele of rs231642830 (AB) and non-carrier (AA) controls. Data are mean with biological replicates shown as individual data points or noted in figure, in F) and G) error bars represent S.E.M. * P < 0.05 compared to AA mice.

To stratify mice for further analysis we conducted single nucleotide polymorphism (SNP) analysis of the defensin locus. We identified one intergenic SNP rs231642830, and one structural variant SV_8_21749161_21749163, with genome-wide significant logarithm of the odds (LOD) scores (Figure 1D). We categorised mice by the number of minor alleles they carried at the rs231642830 SNP (AA = no minor alleles, AB = one copy, BB = two copies). Animals carrying the AB allele had identical glucose tolerance to ‘control’ AA mice despite a 50% reduction in insulin at the 15-minute time point (Figure 1E,F) and increased insulin sensitivity determined by the Matsuda Index (Figure 1G). These data suggest genetic variance at the defensin locus is linked to improved whole body insulin sensitivity.

### A single nucleotide polymorphism within the defensin locus associates with insulin sensitivity and Akkermansia muciniphila abundance

In mice, defensins are secreted from Paneth cells in the intestinal crypt into the gut lumen to modulate microbial composition (Figure 2A). Therefore, we performed 16S rRNA sequencing to profile the microbiome composition of caecal contents of mice from three groups: 1) mice carrying the putative protective rs231642830 allele (AB), 2) cage mates of these mice, and 3) a subset of AA control mice (Figure 2B). Because mice are coprophagic, analysing cage mates allows us to test for co-housing effects on insulin sensitivity, evidence for which would support a microbiome mediated mechanism of action.

**Figure 2.**
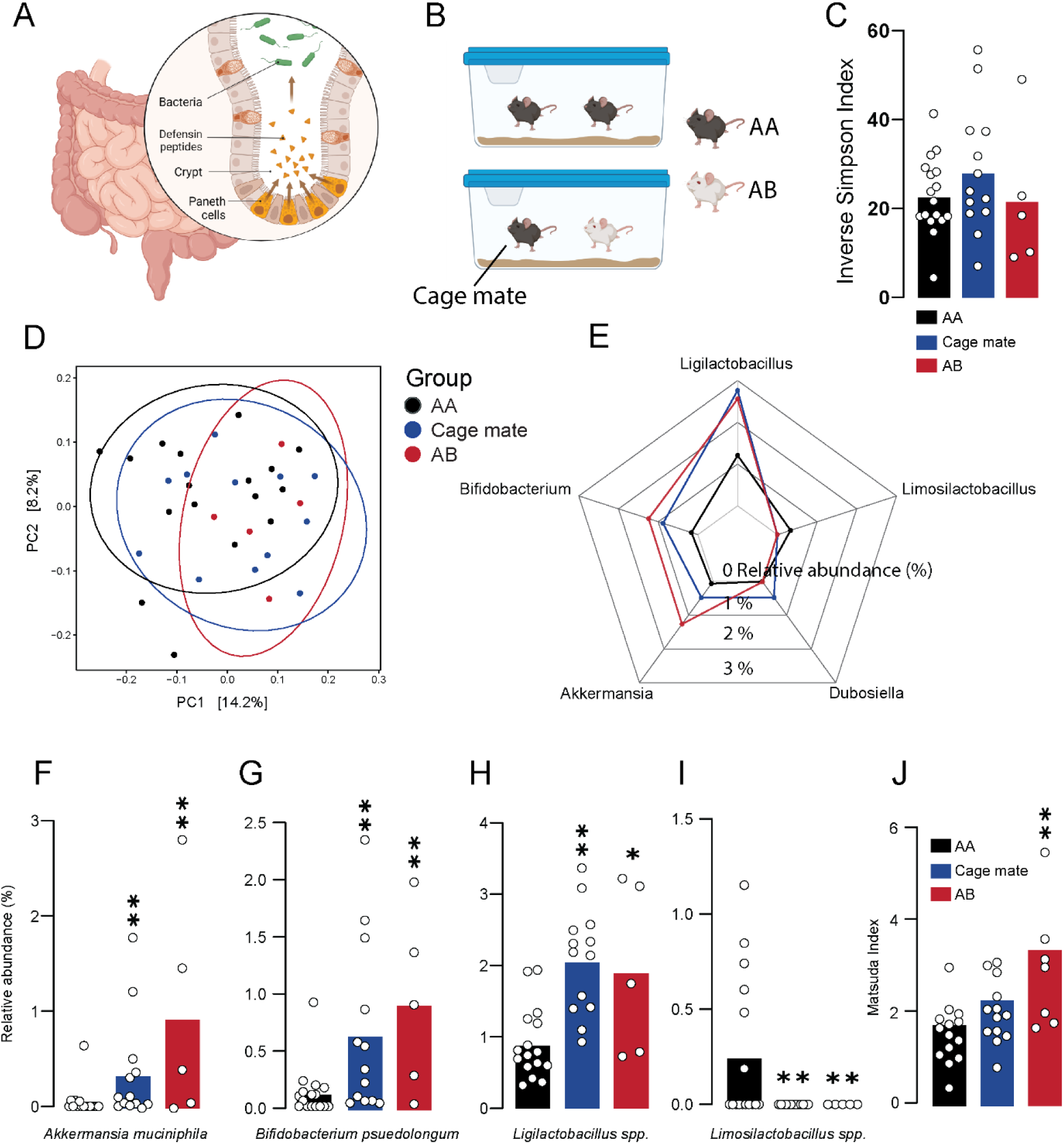
– Analysis of microbial composition and co-housing effects in mice carrying a putative insulin sensitivity allele within the defensin locus. **A)** Schematic of murine defensin secretion from Paneth cells into the small intestine. **B)** Schematic of the study design including mice carrying rs231642830 (AB), non-carrier controls (AA), and the cage mates of mice carrying rs231642830 (Cage mate). **C)** Alpha diversity (Inverse Simpson) of AB, AA and Cage mates. **D)** Visualisation of beta-diversity between AB, AA and Cage mate mice calculated by Bray-Curtis dissimilarity. **E)** Relative abundances of differentially represented microbes between AB, AA and Cage mate mice. **F)** Relative abundance of *Akkermansia muciniphila* in AB, AA and Cage mate mice. **G)** Relative abundance of *Bifidobacterium psuedolongum* in AB, AA and Cage mate mice. **H)** Relative abundance of *Ligilactobacillus spp*. in AB, AA and Cage mate mice. **I)** Relative abundance of *Limosilactobacillus spp*. in AB, AA and Cage mate mice. **J)** Matsuda Index in AB, AA and Cage mate mice. Data are mean with biological replicates shown as individual data points. ** P < 0.01, * P < 0.05 compared to AA mice.

All AB mice had been separately housed, and AA control mice were chosen at random from across 7 different cages that had no AB mice. No significant difference in alpha diversity (Inverse Simpson) was observed between groups (Figure 2C). However, beta-diversity analysis (Bray-Curtis dissimilarity and PERMANOVA) (Figure 2D) of the three groups revealed a trend towards differences in overall microbiome composition between AB mice and AA controls (p value = 0.05), and between AA controls and cage mates of AB mice (p value = 0.055), but not between AB mice and their cage mates (p value = 0.622). Analysis of the combined groups (AB mice + cage mates), against the AA control mice revealed divergent microbiomes (p value = 0.009).

Differential abundance analysis of these groups by Analysis of Compositions of Microbiomes with Bias Correction (ANCOM-BC) revealed that compared to AA controls, *Akkermansia muciniphila* (Figure 2E, F), *Bifidobacterium pseudolongum* (Figure 2E, G) and *Ligilactobacillus spp.* (Figure 2E, H), were higher in cage mate and AB mice, while *Limosilactobacilius spp*. (Figure 2E, I) were lower. Intriguingly, *A. muciniphila* has strong links to metabolic health and has been shown to increase in response to α-defensin administration in C57BL/6J mice (11, 12, 29, 43, 44). These data are consistent with AB mice possessing altered microbiomes relative to AA mice, and this can be transmitted to their cage mates. Critically, cage mates of AB mice also trended towards greater insulin sensitivity relative to AA control mice (Figure 2J) suggesting that insulin sensitivity differences between AA and AB mice may be transferable by cohousing via coprophagy.

### Alpha defensin 26 positively correlates with insulin sensitivity and with founder strain contributions towards the Matsuda QTL in the defensin locus

We next sought to identify the specific defensin isoform that confers increased insulin sensitivity. To do this we quantified small intestine defensin isoform protein expression and insulin sensitivity in the Diversity Outbred founder strains (Figure 3A). One advantage of genetic analyses in DOz is that the QTL analysis also provides the contribution of each founder strain towards the QTL signal, and this allows validation experiments to be conducted in the founder strains. Consistent with our previous work (30) and that of others (33, 34), we observed significant variation in glucose tolerance (Figure S1A-H), insulin sensitivity, and adiposity between the inbred founder strains (Figure 3B-C). Using liquid chromatography coupled to tandem mass spectrometry we detected 9 distinct defensin isoforms across 7 strains. To rule out differences in Paneth cell abundance (25) influencing the apparent defensin expression levels, we normalised defensin peptide expression to the average of all defensins within each strain. Correlation analysis of normalised defensin isoform expression revealed that a single isoform, α-defensin 26 (Defa26), was positively correlated with insulin sensitivity (R = 0.73, P < 0.05; Figure 3D,E). We then calculated the QTL founder effects for Matsuda Index on the chromosome 8 locus: A/J, NOD/ShiLtJ, and to a lesser extent 129S1/SvlmJ, contributed positively while CAST/EiJ, NZO/HILtJ, WSB/EiJ, C57BL/6J and PWK/PhJ contributed negatively towards the QTL (Figure 3F). Correlating the founder effects at 21.7 MBp on chromosome 8 with mean normalised defensin levels revealed that only alpha-defensin 26 levels varied in accordance with the contributions of each founder strain to the QTL (R = 0.89, P < 0.01; Figure 3G).

**Figure 3.**
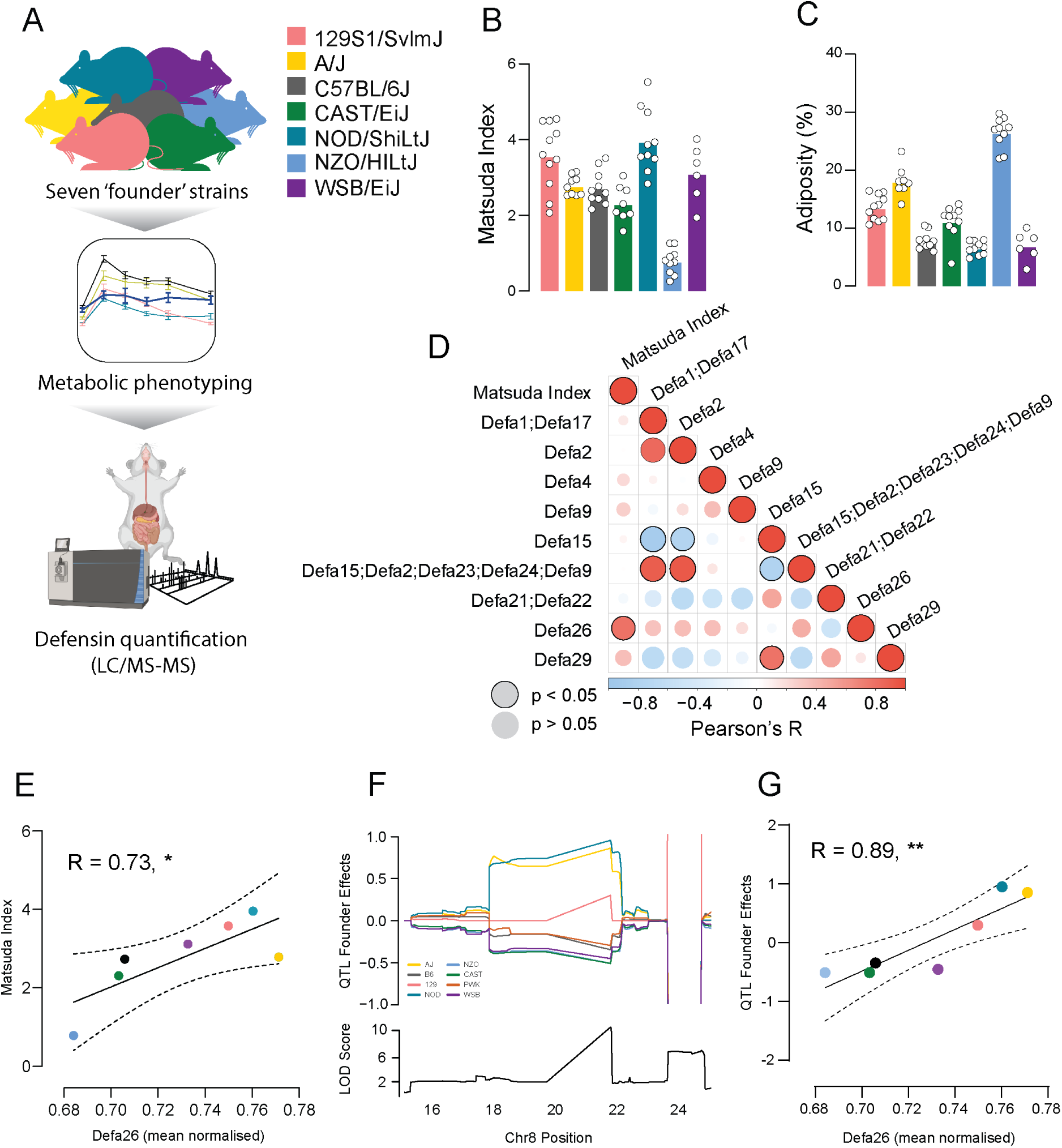
– Analysis of defensin protein expression in Diversity Outbred founder strains. **A)** Schematic of study design to investigate insulin sensitivity and defensin protein expression in small intestines of inbred mouse strains. **B)** Matsuda Index in Diversity Outbred founder strains. **C)** Adiposity in Diversity Outbred founder strains. **D)** Correlations of all quantified defensin peptides (Defa) with Matsuda Index across Diversity Outbred founder strains. **E)** Correlation of mean normalised alpha-defensin 26 abundances with Matsuda Index in Diversity Outbred founder strains. **F)** Founder strain contribution estimates for the DOz Matsuda Index QTL (top) and QTL LOD score (bottom) on chromosome eight. **G)** Correlation of founder strain contribution estimates for the DOz Matsuda Index QTL with mean normalised alpha-defensin 26 abundance. Dashed lines denote 95% confidence intervals. Data are mean with biological replicates shown as individual data points. ** P < 0.01, * P < 0.05

### Alpha defensin 26 dietary supplementation improves insulin sensitivity in HFD-fed C57BL/6J mice

Genetic mapping in DOz mice and analysis of the founder strains revealed that alpha-defensin 26 is a positive regulator of insulin sensitivity. To explore whether alpha-defensin 26 could protect against diet-induced insulin resistance we undertook dietary supplementation studies by synthesising the lumenal (to mimic post-processing secretion) form of alpha-defensin 26 by 9-fluorenylmethyloxycarbonyl-solid-phase peptide synthesis (Fmoc-SPPS), followed by folding (45). As a positive control, we synthesised the lumenal form of human alpha-defensin 5 as this peptide has previously been shown to improve glucoregulatory control in C57BL/6J mice (29). Consistent with previous work (29), mice fed a western diet (WD) supplemented with Fmoc-SPPS synthesised alpha defensin 5 (Defa5) had attenuated weight gain and reduced adiposity but normal lean mass (Figure S2A-C) when compared to control mice fed a control WD. Defa5 supplementation also improved insulin sensitivity, evidenced by equivalent glucose tolerance but lower insulin levels relative to control mice (Figure S2D, E). These results indicate that Fmoc-SPPS synthesised alpha-defensin peptides behave comparably to previous peptides generated by traditional expression systems (26, 27, 29). Based on these results we proceeded with alpha-defensin 26 supplementation (Defa26).

Male C57BL/6J mice were fed either a control WD or WD supplemented with alpha-defensin 26 (WD + Defa26) for eight weeks. On average WD + Defa26 mice gained less overall body mass and adipose tissue (Figure 4A, B), but had equivalent lean mass relative to control WD fed mice (Figure 4C). This reduction in adipose tissue does not appear to be the result of reduced food intake as this was comparable between groups (Figure 4D). Although both groups exhibited near identical glucose tolerance (Figure 4E), WD + Defa26 fed mice had lower circulating levels of insulin at the 15 min timepoint of a GTT (Figure 4F) suggesting improved insulin sensitivity. Consistent with this, the Matsuda Index was higher in WD + Defa26 fed mice relative to WD fed controls.

**Figure 4.**
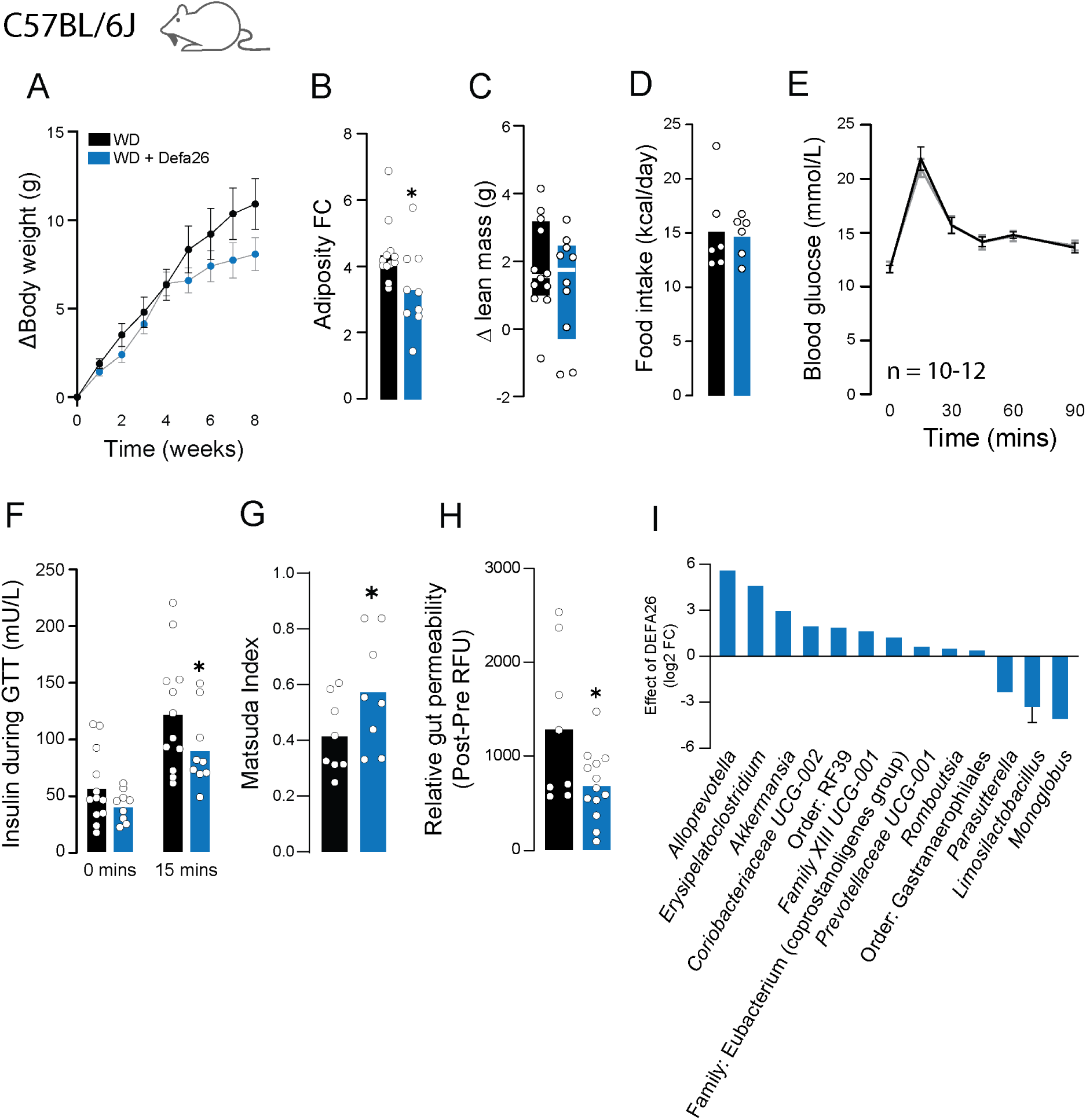
– Metabolic phenotyping of western-diet fed C57BL/6J mice supplemented with alpha-defensin 26. **A)** Change in body weight (g) of western diet (WD) or WD + alpha-defensin 26 (WD + Defa26) fed C57BL/6J mice over eight weeks. **B)** Relative (fold-change) increase in adipose tissue mass of WD and WD + Defa26 fed C57BL/6J mice over eight weeks. **C)** Change in lean mass (g) of WD and WD + Defa26 fed C57BL/6J mice over eight weeks. **D)** Food intake (kcal/day) of WD and WD + Defa26 fed C57BL/6J mice over eight weeks. **E)** Blood glucose concentrations during a glucose tolerance test of C57BL/6J mice fed either a WD or WD + Defa26 for eight weeks. **F)** Blood insulin concentrations during a glucose tolerance test of C57BL/6J mice fed either a WD or WD + Defa26 for eight weeks. **G)** Matsuda Index of C57BL/6J mice fed either a WD or WD + Defa26 for eight weeks. **H)** Relative gut permeability (post – pre FITC fluorescence) of C57BL/6J mice fed either a WD or WD + Defa26 for eight weeks. **I)** Difference (log2 FC) in relative abundance of differential abundant microbes between C57BL/6J mice fed either a WD or WD + Defa26 for eight weeks. Data are mean with biological replicates shown as individual data points. For differentially abundant microbes, error bars represent SD of difference between groups. * P < 0.05 denotes statistical significance from WD control.

In an attempt to profile potential mechanisms underpinning improved insulin sensitivity in WD + Defa26 mice we measured gut permeability by measuring FITC fluorescence in plasma, following oral gavage of FITC-dextran. Consistent with reduced gut permeability, WD + Defa26 fed mice had lower fluorescence than WD fed controls. Considering that genetic variance in the defensin locus associated with increased abundance of certain metabolically beneficial microbes we carried out 16S rRNA sequencing of caecal contents from WD and WD + Defa26 fed mice. In validation of associations between defensin locus SNPs and microbial taxa, caecums from WD + Defa26 fed mice were enriched for *A. muciniphila* and depleted of *Limosilactobaccilus spp*. We also detected increased *Alloprevotella spp.* a microbe whose abundance has previously been linked to defensin supplementation (29).

These results suggested that Defa26 supplementation could protect against WD induced insulin resistance potentially via reduced adiposity and improved gut barrier integrity. Furthermore, caecum microbial composition of WD + Defa26 fed mice exhibits similar changes than the ones observed in mice harbouring a putative protective SNP within the defensin locus.

### Alpha defensin 26 dietary supplementation induces hypoinsulinemia, glucose intolerance and muscle wasting in WD-fed A/J mice

In view of the responses observed in C57BL/6J mice, we next performed experiments A/J mice. A/J mice are protected from diet-induced insulin resistance and express relatively high levels of Defa26 and so we hypothesised that dietary supplementation in this strain would have no effect on whole-body metabolism. Unlike in C57BL/6J mice, A/J mice fed WD + Defa26 exhibited comparable weight gain to WD fed controls (Figure 5A). Furthermore, adiposity increased to the same extent in both WD and WD + Defa26 fed A/J mice (Figure 5B). However, A/J mice fed a WD + Defa26 exhibited a striking (∼1g) reduction in lean mass relative to WD fed controls (Figure 5C). This decrease appears to be the result of muscle wasting, based on summed weights of gastrocnemius, tibialis anterior and quadriceps muscles from WD and WD + Defa26 animals (Figure 5D). As with C57BL/6J mice, we did not observe a difference in food intake between diets (Figure 5E).

**Figure 5.**
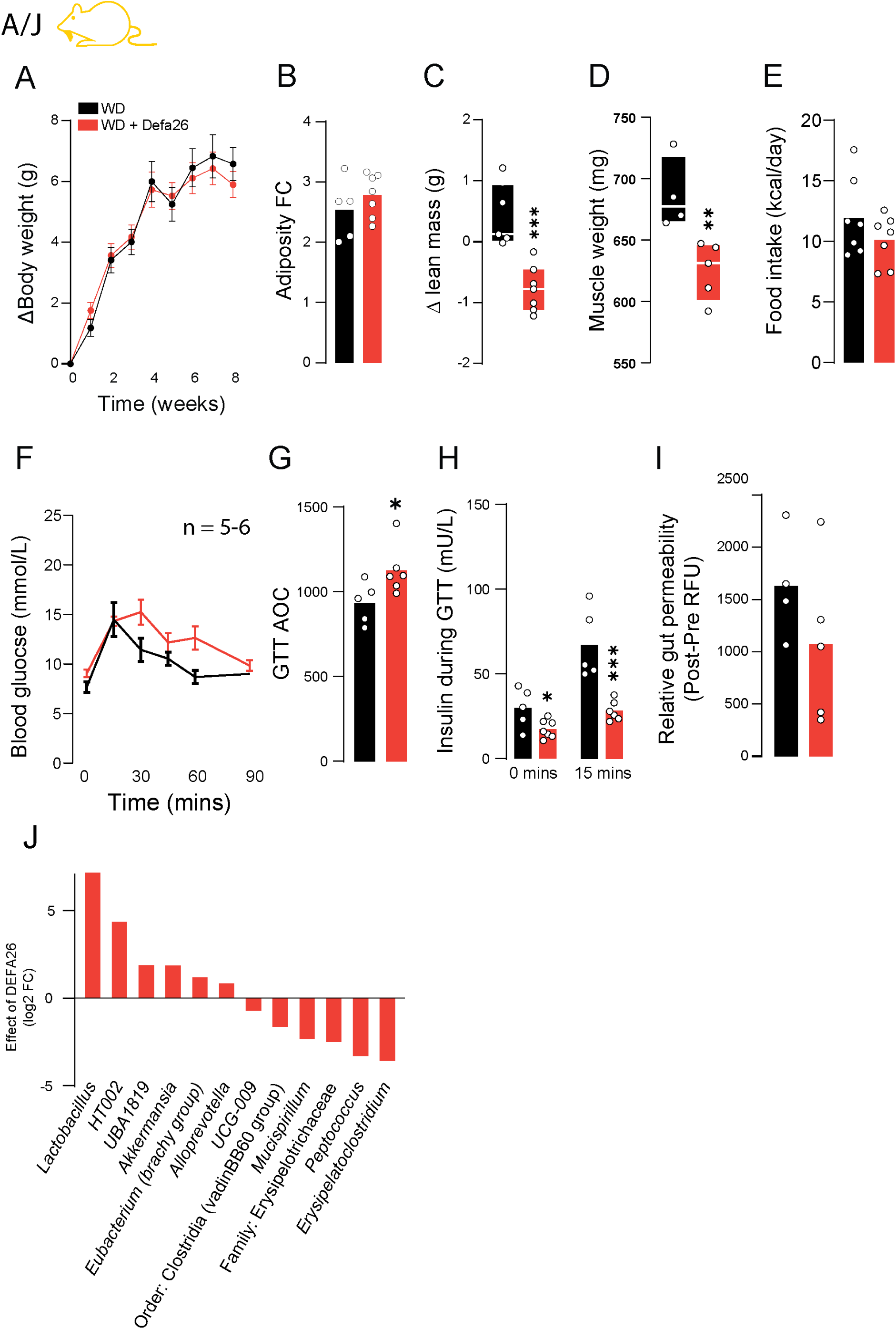
– Metabolic phenotyping of western-diet fed A/J mice supplemented with alpha-defensin 26. **A)** Change in body weight (g) of western diet (WD) or WD + alpha-defensin 26 (WD + Defa26) fed A/J mice over eight weeks. **B)** Relative (fold-change) increase in adipose tissue mass of WD and WD + Defa26 fed A/J mice over eight weeks. **C)** Change in lean mass (g) of WD and WD + Defa26 fed A/J mice over eight weeks. **D)** Combined mass of gastrocnemius, tibialis anterior and quadriceps muscles from A/J mice after WD or WD + Defa26 feeding for 8 weeks. **E)** Food intake (kcal/day) of WD and WD + Defa26 fed A/J mice. **F)** Blood glucose concentrations during a glucose tolerance test of A/J mice fed either a WD or WD + Defa26 for eight weeks. **G)** Glucose tolerance test ‘area-under-the-curve’ for A/J mice fed either a WD or WD + Defa26 for eight weeks. **H)** Blood insulin concentrations during a glucose tolerance test of A/J mice fed either a WD or WD + Defa26 for eight weeks. **I)** Relative gut permeability (post – pre FITC fluorescence) of A/J mice fed either a WD or WD + Defa26 for eight weeks. **J)** Difference (log2 FC) in relative abundance of differential abundant microbes between A/J mice fed either a WD or WD + Defa26 for eight weeks. Data are mean with biological replicates shown as individual data points. *** P < 0.001, ** P < 0.01, * P < 0.05 denotes statistical significance from WD control.

To assess the effect of Defa26 on glucose homeostasis in A/J mice we performed GTTs and observed relative fasting hyperglycaemia and glucose intolerance in WD + Defa26 fed (Figure 5F,G). This appeared to be the result of hypoinsulinemia rather than insulin resistance as WD + Defa26 fed A/J exhibited lower circulating insulin levels both in fasting conditions and during a GTT. Unlike C57BL/6J mice, A/J mice fed WD + Defa26 did not exhibit improvements in gut integrity over WD fed controls (Figure 5I). Despite these differences in phenotypic response relative to C57BL/6J mice, *A. muciniphila* and *Alloprevotella spp* were also enriched in the caecums of A/J mice supplemented with Defa26, albeit to a lesser extent (Figure 5J).

### Disrupted microbial bile acid metabolism may explain the deleterious effects of alpha-defensin 26 supplementation in A/J mice

Despite our hypothesis that Defa26 supplementation would have little to no impact on A/J mice, we in fact observed muscle wasting, glucose intolerance and hypoinsulinemia in response to 8 weeks of dietary supplementation in this strain. This contrasted sharply with the beneficial effects observed in C57BL/6J mice and reinforces the importance of strain selection when testing potential therapeutics.

To begin understanding what might underpin these effects we looked to the microbiome as this is the primary site of action for defensin peptides. Comparing the differentially abundant microbes in C57BL/6J and A/J mice revealed relatively strain-specific changes (Figure 6A,B). Only three microbes were changing concordantly in both strains: *A. muciniphila* (Figure 6C)*, Alloprevotella spp.* (Figure 6D), and *Erysipelatoclostridium spp* (Figure 6E). Further reinforcing strain specificity, *Erysipelatoclostridium spp* increased in C57BL/6J but was depleted by Defa26 treatment in A/J mice (Figure 6E).

**Figure 6.**
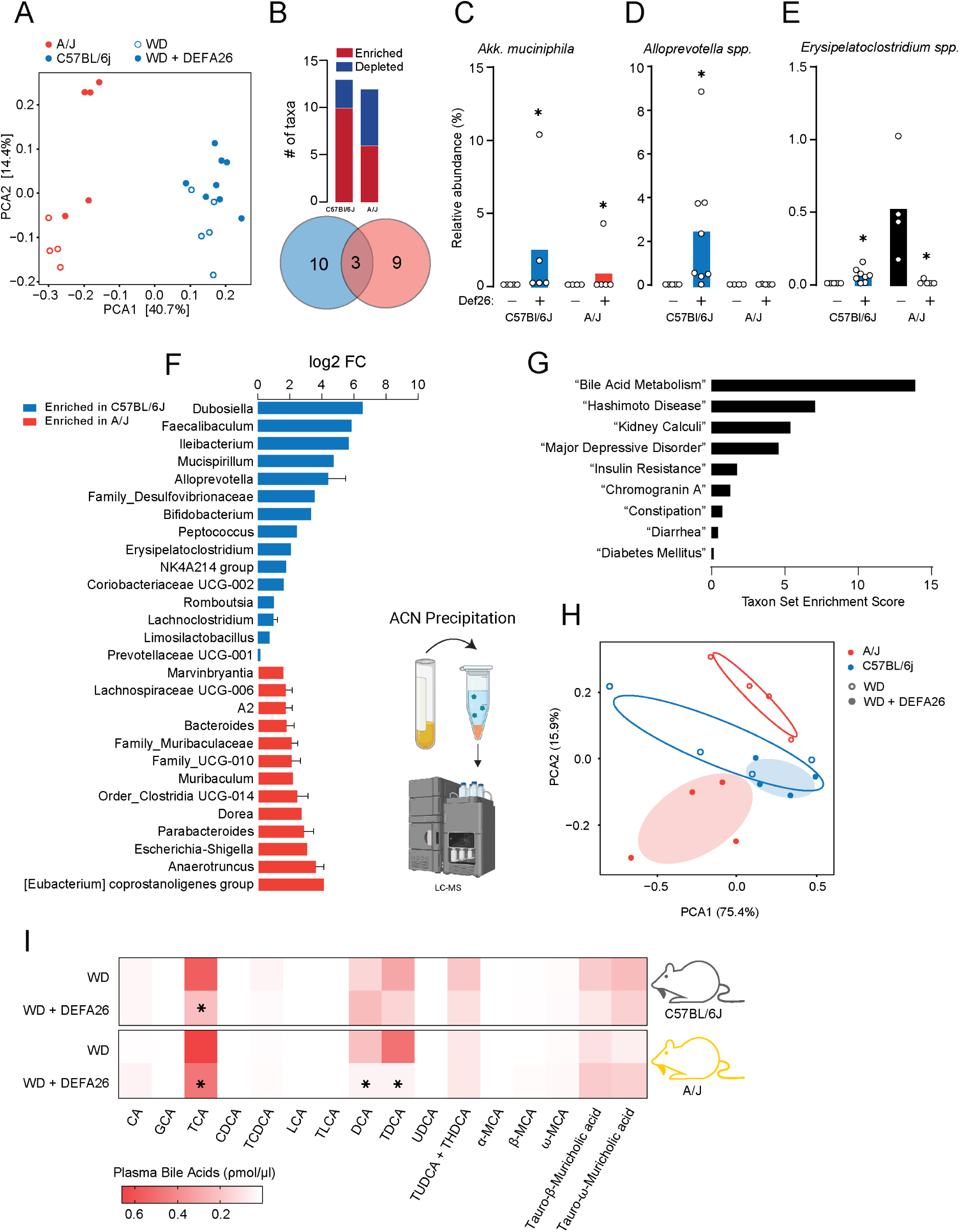
– Comparison of caecal microbiomes and circulating bile acids from C57BL/6J and A/J following alpha-defensin 26 supplementation. **A)** Visualisation of beta-diversity (ANCOM-BC) in WD and WD + Defa26 fed C57BL/6J and A/J mice. **B)** Comparison of differentially abundant microbial taxa in mice fed either a WD or a WD + Defa26 assessed by ANCOM-BC. C) *Akkermansia muciniphila* relative abundance in C57BL/6J and A/J mice fed either a WD or a WD + Defa26. D) *Alloprevotella sp.* relative abundance in C57BL/6J and A/J mice fed either a WD or a WD + Defa26. E) *Erysipelatoclostridium spp.* relative abundance in C57BL/6J and A/J mice fed either a WD or a WD + Defa26. **F)** Difference (log2 FC) in relative abundance of differential abundant microbes between C57BL6/J and A/J mice fed WD + Defa26 for eight weeks. **G)** Enrichment scores for statistically significant terms included in Taxon Set Enrichment Analysis. **H)** Principal component visualisation of circulating bile acids in C57BL/6J and A/J mice fed either a WD or a WD + Defa26. **I)** Heatmap visualisation of circulating bile acid concentrations in C57BL/6J and A/J mice fed either a WD or a WD + Defa26. Data are mean with biological replicates shown as individual data points. * P < 0.05 denotes statistical significance from WD control.

To further investigate how microbiome-strain interactions could underpin the differential effects of Defa26 supplementation we compared the microbiomes of Defa26 treated C57BL/6 and A/J mice (Figure 6F). This revealed 28 differentially abundant taxa, which were then analysed by taxon-set enrichment analysis (46). Using the ‘host-intrinsic’ dataset revealed a striking enrichment for ‘Bile Acid Metabolism’, as well as other relevant terms including ‘Insulin Resistance’ and ‘Diabetes Mellitus’ (Figure 6G). Based on this, and previous work linking microbially derived bile acids to insulin secretion and lean mass (16, 47, 48) we set out to profile circulating bile acids in Defa26 fed mice.

Using liquid chromatography-mass spectrometry (LC-MS) we measured the abundance of 16 primary and secondary bile acids in plasma from a subset of C57BL/6J and A/J mice fed either a WD or WD + Defa26 (S3A, B). Consistent with strains-specific effects of Defa26 supplementation, principal component analysis could separate A/J but not C57BL/6J bile acid profiles based on Defa26 supplementation (Figure 6H). In A/J mice, Defa26 has a mild effect on taurocholic acid (TCA) levels, as well as more striking reductions in deoxycholic acid (DCA) and taurodeoxycholic acid (TDCA) (Figure 6I, S3B). Notably, both DCA and TDCA are derived from microbially mediated deconjugation reactions and can activate the canonical bile acid receptor FXR (farnesoid X receptor) (49). Previous work has shown that activation of FXR in pancreatic beta-cells by bile acids is required for optimal glucose-stimulated insulin secretion (48, 50) and bile-acid induced FXR signalling in the liver can modulate muscle mass via FGF15/19 (17). Therefore, reductions in DCA and TDCA in A/J mice due to disrupted microbial bile acid metabolism may explain the observed effects on reduced insulin secretion and muscle mass.

## Discussion

Functional links between the gut microbiome and glucose homeostasis have predominately been made via high fat/high sugar, low fibre ‘western’ diets, which bias gut microbial composition towards an inflammatory obesogenic state (8, 51, 52). Here we reveal an alternative gut-microbiome/metabolic health axis by performing genetic mapping in a population of chow-fed DOz mice. We identified a striking insulin sensitivity QTL within the defensin gene cluster, which associated with an enrichment for metabolically beneficial microbes, and enhanced expression of the antimicrobial peptide Defa26. We validated this observation by performing dietary supplementation studies in two inbred mouse strains with differential endogenous Defa26 expression. Our results revealed that Defa26 controls glucose homeostasis in an inverted U-shaped relationship. At low levels, increasing concentrations of Defa26 improved gut integrity and *A. muciniphila* abundance, whereas excess Defa26 disrupted microbial bile acid metabolism, leading to insulin secretion defects and muscle wasting. This illustrates the importance of considering genetic variation in the development of metabolic therapeutics and places the microbiome downstream of host genetics in the control of insulin sensitivity.

In our DOz colony, litters are separated at weaning to avoid confounding genetic diversity with cage-effects. We took advantage of this to test whether the defensin locus/insulin sensitivity association was microbiome mediated. We observed that sharing a cage with an AB mouse conferred a mild beneficial effect on insulin sensitivity and increased *A. muciniphila* abundance. This is consistent with previous defensin supplementation studies (43, 53) and microbial transfer via coprophagy, and suggests the mechanism linking the defensin locus to insulin sensitivity is microbiome mediated. This has important implications beyond the present study as siblings of different genotypes are commonly co-housed in traditional transgenic mouse experiments. If there is an effect of genotype on gut microbial composition, any subsequent effect on host-physiology may be masked by microbial transfer between cage-mates. For example, a potential Defa26 knock-out mouse model co-housed with wild-type control mice may appear phenotypically normal provided it maintains a healthy microbiome via coprophagy.

The murine defensin cluster on chromosome 8 contains as many as 75 defensin genes, 12 of which were located within 2 Mb of the insulin sensitivity Matsuda Index QTL. To determine which of these were potentially mediating the association between the defensin locus and insulin sensitivity we performed tandem mass spectrometry on small intestine tissue in DOz founder strains. Taking this approach, only Defa26 associated with insulin sensitivity and QTL founder contributions suggesting that genetic variants within the defensin locus promote both Defa26 expression and insulin sensitivity. Defa26 was first identified in 2004 by a phylogenetic search of the *Mus musculus* genome (54). Interestingly, the aforementioned study revealed that while the mature defensin peptide sequence varies between isoforms, the signal peptide and pro-peptide are highly conserved. This suggests that selection has optimised the lumenal activity of defensins rather than their pre-secretion processing. Alignment of the mature defensin peptide sequences identified in our study revealed that Defa26 is unique relative to other defensins at the following residues V63G and K72T. Notably the glycine at position 63 is near the N-terminus of the first beta sheet and may alter interactions between the defensin ‘barrel structure’ and microbial membranes, leading to selective antimicrobial activities and the observed effects on insulin sensitivity.

AB DOz mice and C57BL/6J mice supplemented with alpha-defensin 26 were both more insulin sensitive than their relevant controls and their caecum was enriched for the mucin-dwelling microbe *A. muciniphila*. As defensins are the most concentrated in the mucin layer (22), it stands to reason *A. muciniphila* has evolved resistance against defensin peptides that can be exploited by defensin administration for beneficial metabolic effects by enabling *A. muciniphila* growth that may have been prevented by other defensin sensitive microbes (11, 12, 43). Intriguingly, Zhang et al., (21) identified several significant QTL for *A. muciniphila* in high-fat diet (HFD) fed DO mice. While these QTL did not include the defensin locus, they did include the *Atf3* and *Tifa* loci which are involved in Paneth cell differentiation. Differences between our study and that of Zhang et al., likely reflect differences in diet and population genetic architecture, but nevertheless, they both point towards an important role of Paneth cells and defensins in *A. muciniphila* abundance.

In stark contrast with our hypothesis that A/J mice would not respond to Defa26 supplementation they exhibited hypoinsulinemia, muscle wasting and glucose intolerance. Previous experiments in DO founder strains (16) and TSEA comparing the gut microbiomes of Defa26 fed C57BL/6J and A/J mice indicated this could be due to disrupted bile acid metabolism. Indeed, circulating levels of TCA, DCA and TDCA were reduced by Defa26 supplementation in A/J but not C57BL/6J mice. This suggests that Defa26 supplementation may disrupt microbes that facilitate the stepwise deconjugation and 7α-dehydroxylation reactions ultimately required to convert cholic acid (CA) into TDCA. (49, 55, 56). As agonists for the bile acid receptors FXR and Tgr5, loss of DCA and TDCA may inhibit bile acid signalling in peripheral tissues (57), such as beta-cells which require active FXR signalling for optimal insulin secretion (48, 50) and muscle tissue which exhibits atrophy upon Tgr5 ablation. This disruption may not have occurred in C7BL6/J mice as they exhibit lower endogenous Defa26 expression which ultimately does not reach toxic levels following exogenous supplementation.

There are two conclusions we draw from the present study. First, the gut microbiome is a downstream effector of genetic variants which regulate insulin sensitivity. In our data, microbial and metabolic differences between mice fed an identical diet can be explained by genetic variance at a single locus. Historically, determining cause-and-effect between microbes and insulin resistance has been challenging as both are altered by diet. However, by anchoring upon genetics we can infer causality from genotype to phenotype via the proteome and microbiome. Secondly, and perhaps most importantly, the impact of individual biological differences on potential therapeutic outcomes is significant, and must be considered as we move into the era of preclinical precision medicine.

## Methods

### Mouse breeding and phenotyping

Male ‘Diversity Outbred from Oz’ (DOz) mice were bred and housed at the Charles Perkins Centre, University of Sydney, NSW, Australia as previously described (39). The DOz mice used in this study were outbred for 27 to 36 generations and comprised a total of 670 male DOz mice across 9 separate cohorts. Genomic DNA was isolated from each mouse and subjected to SNP genotyping (58), followed by genotyping diagnostics and cleaning as described (59). Experiments were performed in accordance with NHMRC guidelines and under approval of The University of Sydney Animal Ethics Committee, approval numbers #1274 and #1988. To delineate genetic from cage-effects, mice were randomised into cages of 3-5 at weaning. All mice were maintained at 23°C on a 12-hour light/dark cycle (0600–1800) and given *ad libitum* access to a standard laboratory chow diet containing 16% calories from fat, 61% calories from carbohydrates, and 23 % calories from protein or a high-fat high-sugar diet (western diet; WD) containing 45% calories from fat, 36% calories from carbohydrate and 19% calories from protein (3.5%g cellulose, 4.5%g bran, 13%g cornstarch, 21%g sucrose, 16.5%g casein, 3.4%g gelatine, 2.6%g safflower oil, 18.6%g lard, 1.2%g AIN-93 vitamin mix (MP Biomedicals), 4.95%g AIN-93 mineral mix (MP Biomedicals), 0.36%g choline and 0.3%g L-cysteine). Fat and lean mass measures were acquired via EchoMRI-900 (EchoMRI Corporation Pte Ltd, Singapore) at 14 weeks of age. Glucose tolerance was determined by oral glucose tolerance test (GTT) at 14-weeks of age by fasting mice for 6-hours (0700-1300 hrs) before oral gavage of 20% glucose solution in water at 2 mg/kg lean mass. Blood glucose concentrations were measured directly by handheld glucometer (Accu-Chek, Roche Diabetes Care, NSW, Australia) from tail blood 0, 15, 30, 45, 60, 90 minutes after oral gavage of glucose. Blood insulin levels at the 0– and 15-minute time points were measured by mouse insulin ELISA Crystal Chem USA (Elk Grove Village, IL, USA) according to manufacturer instructions. Blood glucose and insulin levels were integrated into a surrogate measure of whole-body insulin sensitivity using the Matsuda Index:

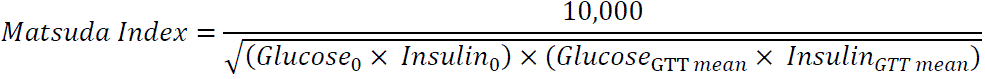

### Genetic mapping analysis

Genetic mapping of Matsuda Index was performed in R using the QTL2 package (60) following square root transformation of raw values. The GIGA-MUGA single nucleotide polymorphism array was used as genomic inputs for mapping (58), and a covariate and kinship matrix to account for genetic relatedness amongst the DOz animals. Significance thresholds were established by performing 1000 permutations and set at *P* < 0.05.

### Caecal DNA isolation

Genomic DNA was extracted from caecal contents of mice using the FastDNA Spin Kit for Feces (MP Biomedicals) as per the manufacturers protocol. DNA concentration was measured using the Qubit dsDNA BR assay kit (Invitrogen). Mock preparations covering all steps of the procedure were conducted as contamination process controls.

### 16S rRNA gene amplicon sequencing analysis

16S rRNA gene amplicon sequencing was performed on all caecal DNA samples. Barcoded amplicon libraries spanning the V4 hypervariable region of the 16S rRNA gene were prepared (515F-806R primer set-515F: GTGYCAGCMGCCGCGGTAA, 806R: GGACTACNVGGGTWTCTAAT) and sequenced using the Illumina MiSeq v2 2 x 250 bp platform at the Ramaciotti Centre for Genomics (UNSW, Sydney, Australia). Raw sequence reads were processed using the DADA2 R package which involves using error profiles to define Amplicon Sequence Variants (ASVs)(61). ASVs were assigned to taxonomy using a pre-trained naïve Bayes classifier trained on the curated 16S rRNA gene SILVA (v138.1) reference database. Any ASV that was present in fewer than 5% of samples or had less than 0.01% of total reads was filtered from the final dataset prior to downstream analysis. Sequencing depth analyses and rarefaction were performed with the *phyloseq* and *vegan* R package (62).

Analysis and graphical presentation of the resultant ASV data was performed in R using the packages *phyloseq*, *vegan*, *microbiome* and *ggplot2*. Alpha diversity metrics were calculated using Inverse Simpson’s index. Beta diversity was assessed on centred-log-ratio transformed ASV counts using Bray Curtis dissimilarity and UniFrac distance and principal coordinate plots generated from the resultant dissimilarity matrix. PERMANOVA (adonis) using the *vegan* R package was used to assess variance in the distance matrices between groups. Differential abundance analysis was performed using the *ANCOM-BC* R package(63).

### Intestinal proteomic sample preparation

Division of the small intestine into thirds was achieved by folding the small intestine into three equivalent lengths and taking a 1 cm section of tissue from the centre of each third. These pieces tissue representing the foregut, midgut and hindgut of each mouse were combined and snap frozen in liquid nitrogen. Frozen samples were then boiled in 400 uL of SDC buffer (4% sodium deoxycholate, 100mM Tris-HCl pH 8.0) by heating at 95 C for 10 minutes at 1000 rpm. Samples were then lysed by sonication for 10 minutes (30 seconds on, 30 seconds off, 70% amplitude protocol). Samples were then heated a second time at 95 °C for 10 minutes at 1,000 rpm before being clarified by centrifugation at 18,000 for 10 minutes at room temperature. Supernatant was taken as lysate and protein concentration was determined by BCA assay, 10 µg of protein was then prepared as previously described (30). Reduction/alkylation (10mM TCEP, 40mM CAA) buffer was added to each sample before incubation for 20 minutes at 60 °C. Once cooled to room temperature, 0.4 ug trypsin and 0.4 ug LysC was added to each sample and incubated overnight (18h) at 37 °C with gentle agitation. 30 µL water and 50 µL 1% TFA in ethyl acetate was added to stop digestion and dissolve any precipitated SDC. Samples were prepared for mass spectrometry analysis by StageTip clean up using SDB-RPS solid phase extraction material (64).(64) Briefly, 2 layers of SDB-RPS material was packed into 200 µL tips and washed by centrifugation at 1,000 x g for 2 minutes with 50 µL acetonitrile followed by 0.2% TFA in 30% methanol and then 0.2% TFA in water. 50 µL of samples were loaded to StageTips by centrifugation at 1,000 g for 3 minutes. Stage tips were washed with subsequent spins at 1,000 g for 3 minutes with 50 µL 1% TFA in ethyl acetate, then 1% TFA in isopropanol, and 0.2% TFA in 5% ACN. Samples were eluted by addition of 60µL 60% ACN with 5% NH_4_OH_4_. Samples were dried by vacuum centrifugation and reconstituted in 30 µL 0.1% TFA in 2% ACN.

### Mass spectrometry analysis

Peptides prepared as above (2 mg total), were directly injected using a Shimazu LC-40 UHPLC onto a 5 cm x 2.1 mm C18 column analytical column (Agilent InfinityLab Poroshell 120 EC-C18, 1.9 um particles) fitted with a 0.5 cm x 2.1 mm C18 guard column (Agilent InfinityLab Poroshell 120 EC-C18, 1.9 um particles). Peptides were resolved over a gradient from 3% – 36% acetonitrile over 10 min with a flow rate of 0.8 mL min^-1^. Peptide ionization by electrospray occurred at 5.5 kV, with curtain gas 25, gas 2 25 and gas 3 35. A 7600 Zeno TOF mass spectrometer (Sciex) with CID fragmentation used for MS/MS acquisition. Spectra were obtained in a data-independent acquisition using Zeno SWATH with 50 isolation width windows spanning 400 to 900 Th. Gas-phase fractionation of a pooled mixture of intestine peptides was performed using 100 Th windows per run, to enable spectral library generation covering the 400-900 Th range. Data files were analyzed using the quantitative DIA proteomics search engine, DIA-NN (version 1.8.1) For spectral library generation, the Uniprot mouse Swissprot database downloaded on the 1st of July 2022 was used. Trypsin was set as the protease allowing for 1 missed cleavage and 1 variable modification. Oxidation of methionine were set as a variable modification. Carbamidomethylation of cystine was set as a fixed modification. Remove likely interferences and match between runs were enabled. Neural network classifier was set to double-pass mode. Protein inference was based on genes. Quantification strategy was set to Robust LC (high accuracy). Cross-run normalization was set to RT-dependent. Library profiling was set to full profiling.

### Peptide synthesis

N,N-dimethylformamide (DMF) and dichloromethane (CH_2_Cl_2_) for peptide synthesis were purchased from RCI Labscan and Merck. Gradient grade acetonitrile (CH_3_CN) for high-performance liquid chromatography was purchased from Sigma Aldrich and ultrapure water (Type 1) was obtained from a Merck Milli-Q EQ 7000 water purification system. Standard Fmoc-protected amino acids (Fmoc-Xaa-OH), coupling reagents and resins were purchased from Mimotopes or Novabiochem. Fmoc-SPPS was performed manually with these reagents and solvents in polypropylene Teflon-fritted syringes purchased from Torviq and through automated synthesis on a SYRO I peptide synthesizer (Biotage). Buffer salts for folding reactions were purchased from Ajax, Sigma Aldrich, and Thermofisher and used as received. Glutathione reduced and oxidised were purchased from Sigma Aldrich and Thermofisher, respectively. All other reagents were purchased from AK Scientific or Merck and used as received.

Electrospray mass spectra (ESI-MS) were obtained using a Shimadzu 2020 UPLC-MS with a Nexera X2 LC-30AD pump, Nexera X2 SPD-M30A UV/Vis diode array detector, and a Shimadzu 2020 mass spectrometer using electrospray ionisation (ESI) operating in positive mode. Separations were conducted using a Waters Acquity UPLC BEH300 (1.7 μm, 2.1 x 50 mm C18 column) with a flow rate of 0.6 mL/min. Spectra are recorded from 300-2000 Da.

Reverse-phase high performance liquid chromatography (RP-HPLC) was carried out on a Waters 2535 Quaternary Gradient system, fitted with a Waters 2489 UV/Vis Detector (monitoring at 214 and 280 nm) and a Waters Fraction Collector III. Linear peptides were purified by preparative RP-HPLC using an Xbridge C18 column (5 μm, 19 × 150 mm) at a flow rate of 12 mL/min. A mobile phase of Milli-Q water (Solvent A) and HPLC-grade CH_3_CN (Solvent B) was employed over a linear gradient with 0.1 vol% TFA (trifluoroacetic acid) as an additive.

Folded peptides were purified by semi-preparative RP-HPLC using a Xbridge C18 column (5 μm, 10 × 250 mm) at a flow rate of 4 mL/min. A mobile phase of Milli-Q water (Solvent A) and HPLC-grade CH_3_CN (Solvent B) was employed over a linear gradient with 0.1 vol% TFA as an additive.

Analytical RP-HPLC was performed on a Waters Alliance e2695 HPLC system equipped with a 2998 PDA detector (λ = 210–400 nm). Separations were performed on a Waters Sunfire® C18 (5 µm, 2.1 × 150 mm) column at 40 °C with a flow rate of 0.5 mL/min. All separations were performed using a mobile phase of 0.1% TFA in water (Solvent A) and 0.1% TFA in CH_3_CN (Solvent B) using linear gradients.

Peptides were synthesized on a 50 µmol scale. 2-Chlorotrityl chloride (2-CTC) resin was treated with Fmoc-Xaa-OH (1.2 eq) and *i*-Pr_2_NEt (4.8 eq) in DCM (4 mL). The C-terminal amino acid of each sequence was used for this loading step, ie. Fmoc-Leu-OH was used for the loading of Defa26 and Fmoc-Arg(Pbf)-OH was used for Defa5. Fmoc loading was determined by measuring the piperidine fulvene adduct from Fmoc deprotection. The loading of the resins were: 0.51 mmol/g for Leu on CTC (Defa26) and 0.50 mmol/g for Arg on CTC (DEFA5).

General procedure A: Automated Fmoc-Solid-Phase Peptide Synthesis (SPPS) –SYRO I automatic peptide synthesizer (Biotage).

50 µmol of the amino acid loaded resin was treated with a solution of piperidine (40 vol%, 0.8 mL) in DMF for 3 min, drained, before repeat treatment with piperidine (20 vol%, 0.8 mL) in DMF for 10 min. The resin was then drained and washed with DMF (4 x 1.2 mL) before addition of a solution of Fmoc-Xaa-OH (200 µmol, 4 eq.) and Oxyma (4.4 eq.) in DMF (400 µL), followed by a solution of *N,N’*-diisopropylcarbodiimide (4 eq.) in DMF (400 µL). The resin was then agitated at 75 °C for 15 min or 50 °C for 30 min as specified [coupling of Fmoc-His(Trt)-OH and Fmoc-Cys(Trt)-OH were reacted at 50 °C for 30 min in all instances]. The resin was then drained via vacuum and one repeat treatment of the coupling conditions was conducted. The resin was then washed with DMF (4 x 1.2 mL) before being treated with a solution of 5 vol% Ac_2_O and 10 vol% *i*-Pr_2_NEt in DMF (1.6 mL) and agitated for 5 min to cap unreacted N-terminal amines on the growing peptide. The resin was then drained and washed with DMF (4 x 1.6 mL). Iterative cycles of this process were repeated until complete peptide elongation was achieved after which the resin was washed with DMF (4 x 5 mL) and CH_2_Cl_2_ (5 x 5 mL). HCl counterion exchanges were performed by dissolving the folded peptide in 0.1 M HCl and lyophilising on a freeze drier. This HCl treatment and lyophilisation was repeated 6 times.

### Synthesis of Defa26

**Scheme 1.**
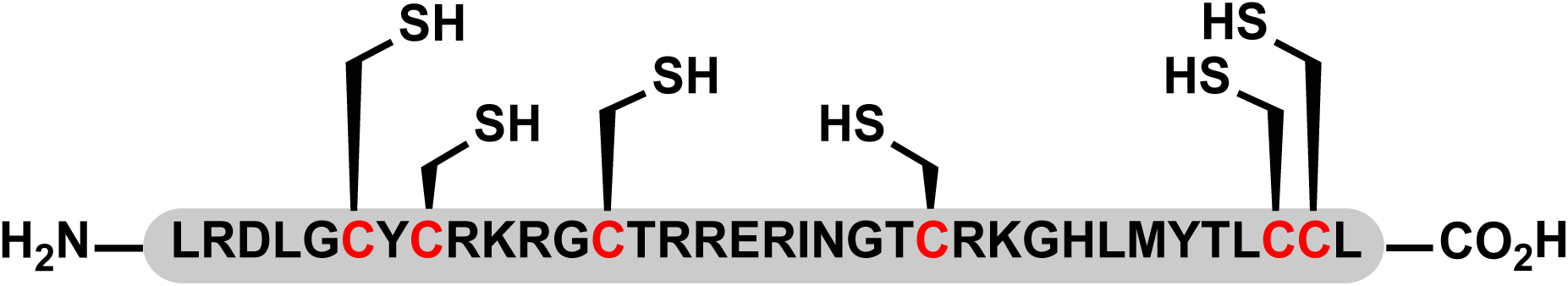
Linear sequence of Defa26.

The above linear sequence was synthesised according to General Procedure A on 2-CTC resin which was loaded with Fmoc-Leu-OH. The peptide was then cleaved from resin by treatment with 85:5:5:5 v/v/v/v TFA/triisopropylsilane/H_2_O/ethanedithiol for 2 h at rt. The cleaved solution was collected, dried to ∼1 mL under N_2_ flow and the peptide product was precipitated using diethyl ether (2 x 40 mL) and collected *via* centrifugation. The crude linear peptide was then purified by preparative RP-HPLC (0 vol% CH_3_CN + 0.1 vol% TFA for 10 min, then 0-50 vol% CH_3_CN + 0.1 vol% TFA over 50 min) and lyophilised affording the linear Defa26 as a white solid (10.14 mg, 4%).

Linear Defa26 (5.5 mg, 1 eq) was first dissolved in 220 µL of rapid dilution buffer containing TRIS (50 mM), NaCl (150 mM), guanidine.HCl (6 M), and tris(2-carboxyethyl)phosphine (2 mM). This solution was then added gradually to a buffer containing NaHCO_3_ (200 mM), urea (2 M), GSH (1 mM), and GSSG (0.2 mM) in MilliQ water to make up a 1 mg/mL peptide solution. The folding reaction was left for 16 h without stirring. The folding progress was monitored through LC-MS, and a loss of 6 Da and a simultaneous retention time shift indicated completion of folding. The folded peptide was then purified by semi-preparative RP-HPLC (0 vol% CH_3_CN + 0.1% TFA for 20 min, then 0-50 vol% CH_3_CN + 0.1 vol% TFA over 50 min), affording the folded Defa26 as a white solid after lyophilization (1.07 mg, 19% isolated yield). Prior to biological assays, the peptide was converted to the HCl salt through a HCl counterion exchange.

**Figure S2.**
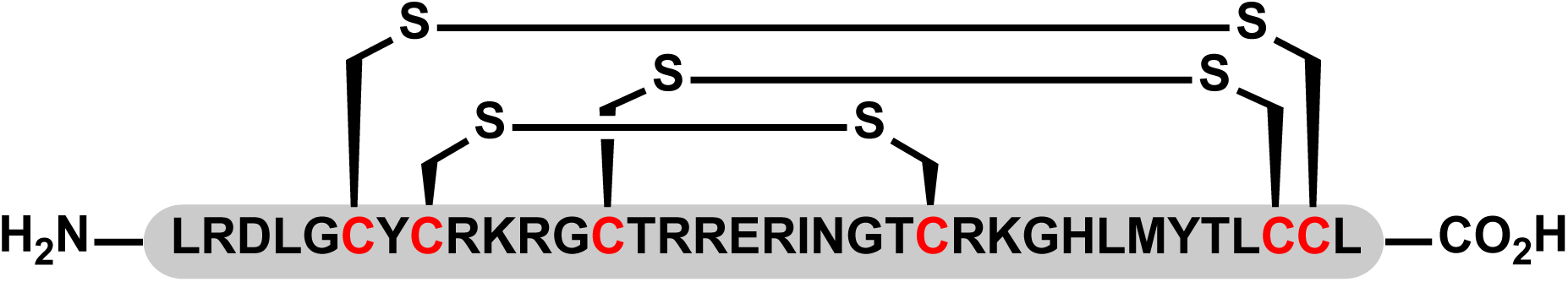
Crude semi-preparative scale trace of folded Defa26. The first 20 minutes of the 0 vol% B + 0.1 % TFA wash is omitted for clarity. Arrow indicates the collected folded peptide.

### Synthesis of DEFA5

**Scheme 3.**
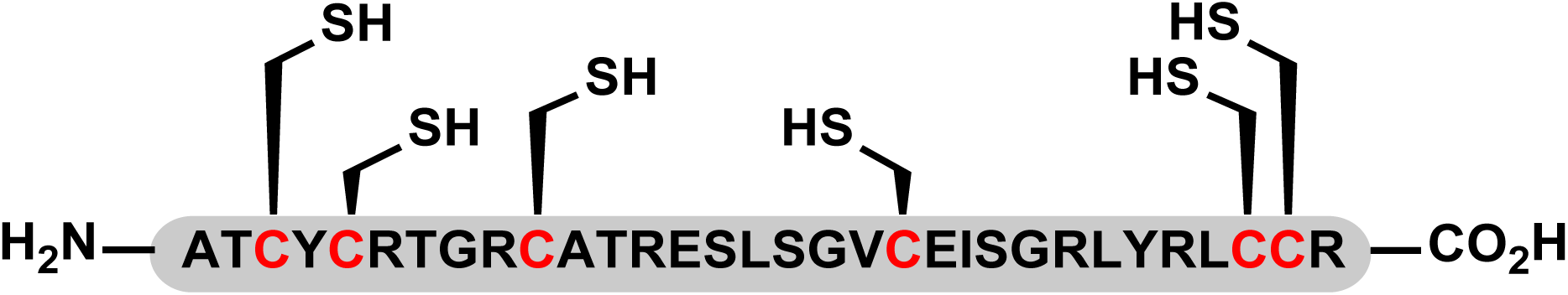
Linear sequence of DEFA5.

The above linear sequence was synthesised according to General Procedure A on 2-CTC resin which was loaded with Fmoc-Arg(Pbf)-OH. The peptide was then cleaved from resin by treatment with 85:5:5:5 v/v/v/v TFA/triisopropylsilane/H_2_O/ethanedithiol for 2 h at rt. The cleaved solution was collected, dried to ∼1 mL under N_2_ flow and the peptide product was precipitated using diethyl ether (2 x 40 mL) and collected *via* centrifugation. The crude linear peptide was then purified by preparative RP-HPLC (0 vol% CH_3_CN + 0.1 vol% TFA for 10 min, then 0-50 vol% CH_3_CN + 0.1 vol% TFA over 50 min), affording the linear DEFA5 as a white solid after lyophilisation (3.19 mg, 3%).

Linear DEFA5 (2.51 mg, 1 eq) was first dissolved in 100 µL of rapid dilution buffer containing TRIS (50 mM), NaCl (150 mM), guanidine.HCl (6 M), and tris(2-carboxyethyl)phosphine (2 mM). This solution was then added gradually to a buffer containing NH_4_OAc (330 mM), guanidine.HCl (500 mM), GSH (1 mM), and GSSG (0.2 mM) in MilliQ water to make up a 1 mg/mL peptide solution. The folding reaction was then left for 40 h without stirring. The folding progress was determined through LC-MS, and a loss of 6 Da and a simultaneous retention time shift indicated folding. The folded peptide was then purified by semi-preparative RP-HPLC (0 vol% CH_3_CN + 0.1% TFA for 20 min, then 0-50 vol% CH_3_CN + 0.1 vol% TFA over 50 min), affording the folded DEFA5 as a white solid after lyophilisation (0.62 mg, 24% isolated yield). Prior to biological assays, the peptide was converted to the HCl salt through a HCl counterion exchange.

**Scheme 4.**
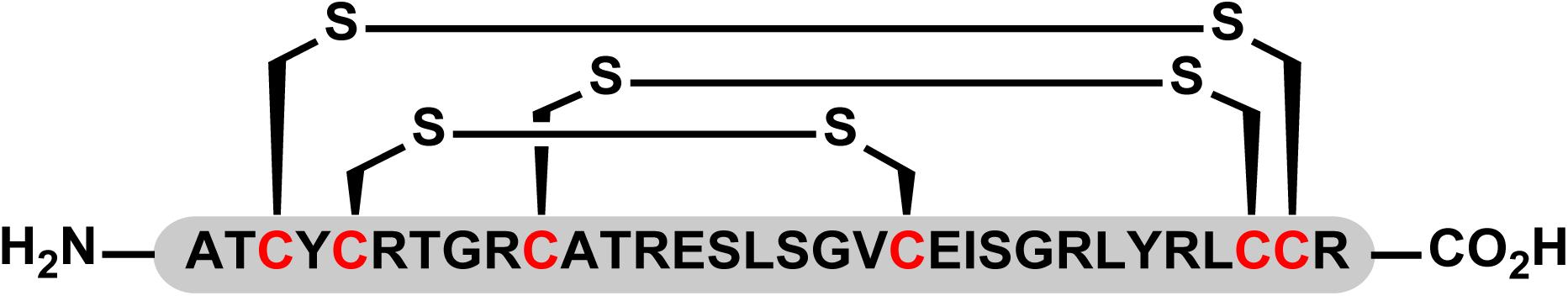
Folded sequence of DEFA5.

### Defensin feeding experiments

Prior to allocation into experimental groups, C57BL/6J and A/J mice underwent baseline metabolic phenotyping was described above. Mice of each strain were then fed a WD or a WD containing the luminal forms of either murine alpha-defensin 26 (Defa26) or human alpha-defensin 5 (DEFA5) for 8 weeks. Synthetic peptides were mixed into mouse diet by hand. Even distribution of peptides in food was monitored by the addition of blue food dye which was used as a proxy for the distribution of peptides throughout each batch. After eight weeks mice underwent a second bout of metabolic phenotyping and assessment of gut permeability via FITC-dextran oral gavage as previously described (65). Briefly mice were fasted for 4 hours (0900–1300) before a baseline blood sample (50 uL) was taken from a tail incision. Mice were then gavaged with 150 μl of 80 mg/ml FITC dextran (4kDa). After 4 hours, a second blood sample was taken. Both samples (baseline and post-gavage were then centrifuged at 5,000 rpm for 10 minutes. Resulting plasma was then diluted 1:10 in PBS and fluorescence was measured at 530 nm with excitation at 485 nm. Data was then expressed as relative fluorescence units.

### Bile acid extraction

50 µL of plasma thawed on ice was added to 150 µL of ice-cold acetonitrile containing 5 pmoles of d4-cholic acid internal standard. Samples were vortexed for 30 s at maximum speed then centrifuged at 15,000 x *g* for 10 min at 4°C to pellet insoluble debris. 170 µL of supernatant was transferred to fused-insert HPLC vials, then vacuum centrifuged to dryness in an Eppendorf Concentrator Plus. Samples were reconstituted in 50 µL of 80:20 water:acetonitrile. All solvents were MS grade.

### Bile acid quantification

Separation of bile acids was performed using a Nexera LC-40 UHPLC (Shimadzu, Rydalmere, NSW, Australia) using a 2.1x 50 mm, 2.7 µm CORTECS C18 column (Waters, Rydalmere, NSW, Australia) with a 7-minute binary gradient of 0.1% formic acid in water (A) and acetonitrile (B) at a flow rate of 0.9 mL/min. Initial gradient conditions of 83:17 A/B rose to 30% B at 1.2 min using curve setting 9. From 1.2 to 3.0 minutes, the proportion of B increased to 38% using curve –5, then rose to 100% at 4.3 min using curve setting 5. The column was flushed at 100% B for 1.9 min before returning to initial conditions over 0.1 min and being held for 0.7 min. Column temperature was 50°C and injection volume was 0.5µL.

MS data were acquired on a ZenoTOF 7600 (Sciex, Mulgrave, VIC, Australia) quadrupole-time-of-flight tandem MS operating with electrospray ionisation in negative polarity. Intact bile acid precursor ions were detected using a TOF MS experiment with mass range 200-600 Da and accumulation time 0.3 s. Source parameters were: Spray voltage: –4500 V, Temperature: 650°C, Ion source gas 1: 70psi, Ion source gas 2: 80psi, Curtain gas: 40psi. Declustering potential was set to –80 V, collision energy was –10 V and CAD gas was set to 10 (arbitrary units). MS calibration was maintained by Calibrant Delivery System auto-calibration at intervals of approximately 1 hour.

Raw data were acquired in a single batch with acquisition order randomised. Six replicates of a sample pool were distributed through the batch to assess intra-batch imprecision. Six replicate injections from a single vial were acquired to determine instrument repeatability. Data analysis was performed with the Analytics module of SCIEX OS (version 3.1.6). Chromatographic peaks were extracted with a width of 0.02 Da and integrated using the AutoPeak algorithm. Identification of bile acids was based on both accurate precursor m/z and retention time matched to commercial standards for all quantified bile acid species (Table 1). Relative molar amounts for bile acids were calculated by comparing raw peak areas relative to the internal standard. Average mass accuracy was >1 ppm, with a range of +/-2ppm across the run. %CVs were calculated as peak area ratios relative to the internal standard.

**Table 1.** Bile acid identification details and coefficients of variation.

| Bile Acid | Mass (Da)<br>[M-H]- | Retention Time<br>(min) | Peak Width<br>(min) | Inter-Batch<br>(%CV) | Intra-Batch<br>(%CV) |
| --- | --- | --- | --- | --- | --- |
| d4 CA | 411.3116 | 2.36 | 0.15 |  |  |
| CA | 407.2803 | 2.36 | 0.11 | 1.17 | 2.15 |
| DCA | 391.2854 | 3.93 | 0.19 | 0.99 | 6.09 |
| CDCA | 391.2854 | 3.74 | 0.16 | 3.63 | 2.21 |
| UDCA | 391.2854 | 2.45 | 0.13 | 5.47 | 4.9 |
| αMCA | 407.2803 | 1.95 | 0.08 | 5.52 | 12.23 |
| βMCA | 407.2803 | 2.01 | 0.09 | 1.81 | 2.45 |
| ωMCA | 407.2803 | 1.91 and 2.01 | 0.08 | 1.93 | 2.53 |
| GCA | 464.3018 | 1.94 | 0.08 | 1.66 | 3.25 |
| LCA | 375.2905 | 4.44 | 0.05 | 13.49 | 25.22 |
| TCA | 514.2844 | 1.71 | 0.10 | 2.17 | 5.32 |
| TDCA | 498.2895 | 2.1 | 0.15 | 1.99 | 6.75 |
| TCDCA | 498.2895 | 2.01 | 0.13 | 2 | 5.05 |
| THDCA | 498.2895 | 1.69 | 0.17 | 2.29 | 4.56 |
| TUDCA | 498.2895 | 1.69 | 0.17 | 2.29 | 4.56 |
| TLCA | 482.2946 | 2.88 | 0.29 | 1.93 | 10.3 |
| TβMCA | 514.2844 | 1.53 | 0.09 | 2.59 | 5.47 |
| TωMCA | 514.2844 | 1.5 | 0.07 | 3.65 | 5.89 |

### Data analysis

All data analysis and visualisation were performed in either the R programming environment (66) or GraphPad Prism (GraphPad Software, San Diego, California USA). For protein correlation analysis the Matsuda Index was calculated using glucose tolerance data before being log2 transformed. To correct for multiple testing, p-values were adjusted using the q-value method in the R package *qvalue* (67).(67) Chi-square tests for distribution differences within the data and two/one-way ANOVA tests for group differences were performed in GraphPad Prism.

## Acknowledgements

We’d like to thank the Sydney Mass Spectrometry facility and Large Animal Services in the Charles Perkins Centre at the University of Sydney for mass spectrometry and mouse housing support.

## Supplementary Figures

**Supplementary Figure 1.**
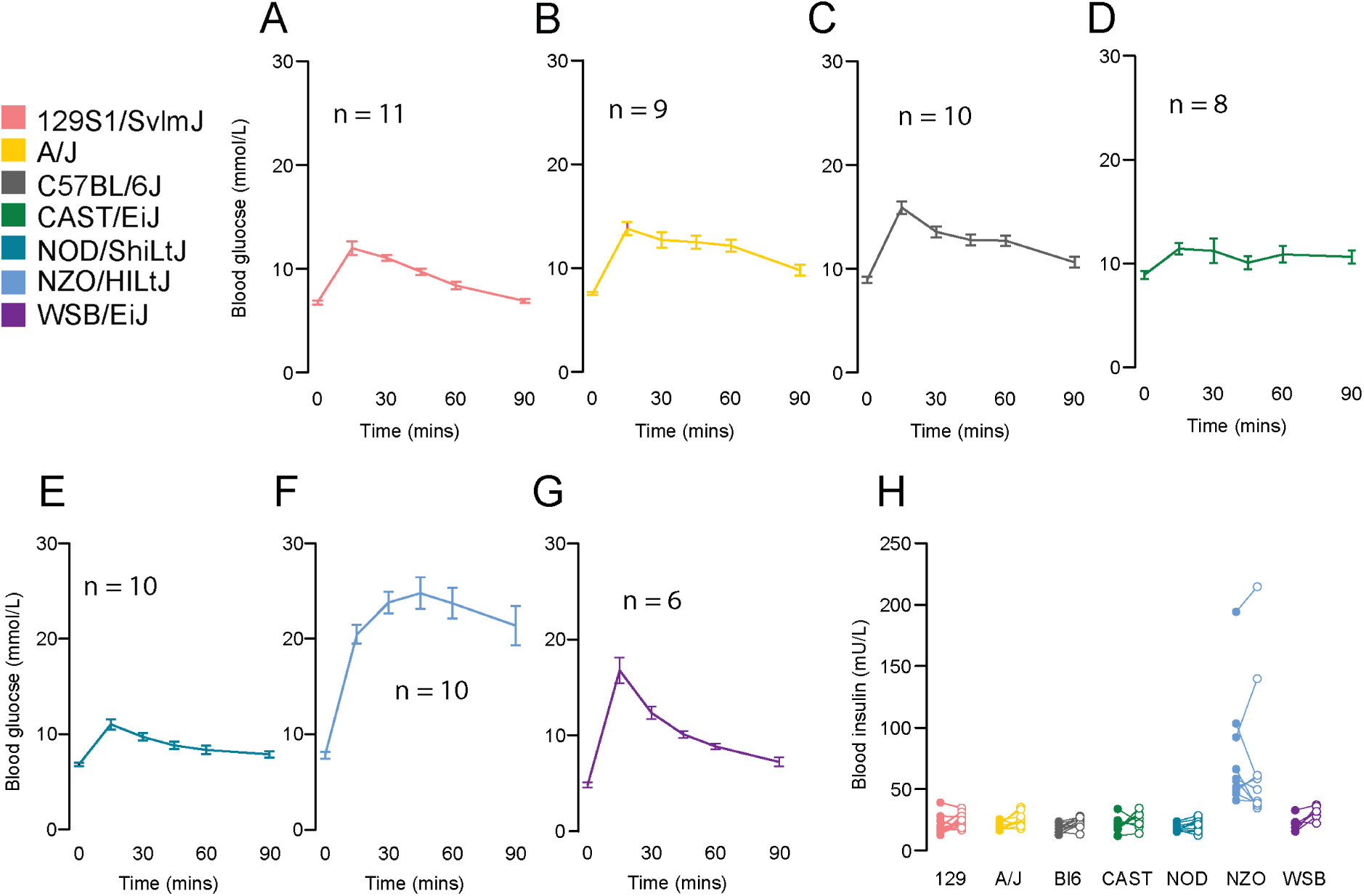
– Glucose tolerance and blood insulin concentrations in Diversity Outbred founder strains. Blood glucose concentrations during a glucose tolerance test in **A)** 129S1/SvlmJ, **B)** A/J, **C)** C57BL/6J, **D)** CAST/EiJ, **E)** NOD/ShiLtJ, **F)** NZO/HILtJ, **G)** WSB/EiJ. **H)** Insulin concentration in Diversity Outbred founder strains during a glucose tolerance test. Data are mean with biological replicates are shown as individual data points or noted in figure. **** P < 0.0001, *** P < 0.001, ** P < 0.01, * P < 0.05

**Supplementary Figure 2.**
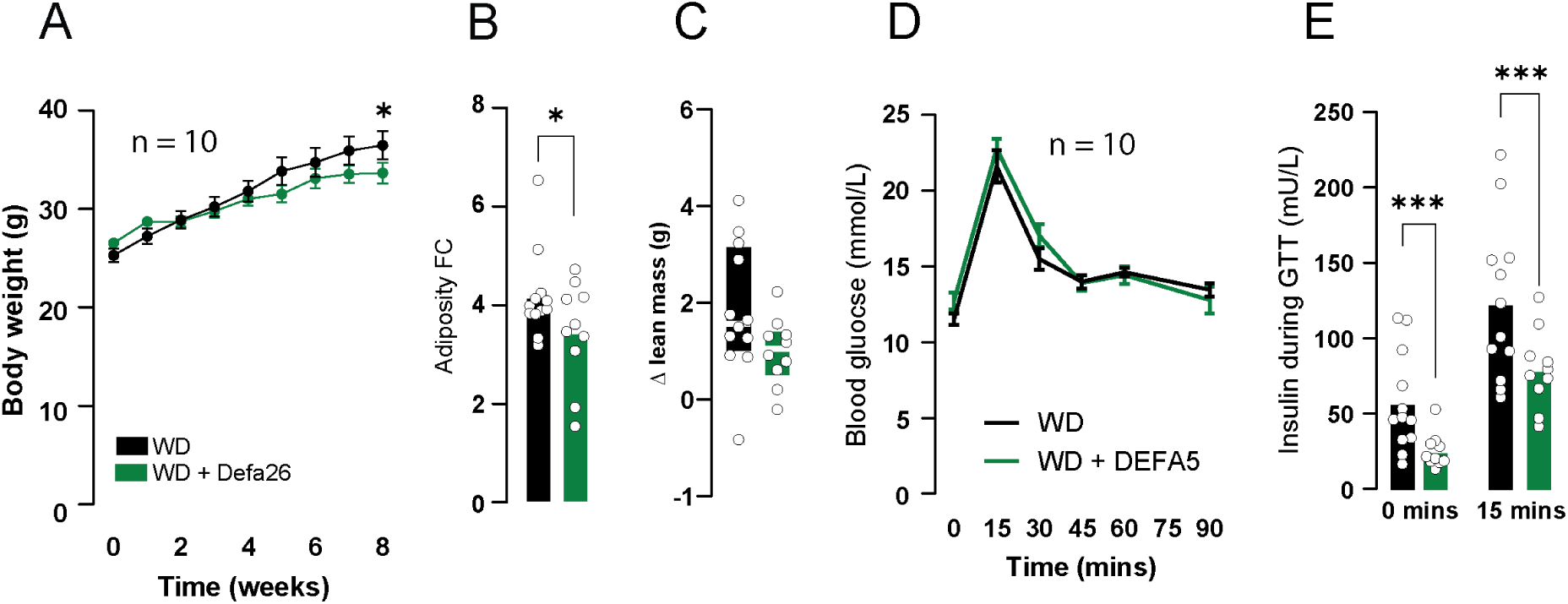
– Effect of alpha-defensin 5 supplementation on insulin sensitivity and body composition in C57BL/6J mice. **A)** Body weight of mice fed either WD or WD+DEFA5. **B)** Fold-change in adiposity in mice fed either WD or WD+DEFA5. **C)** Change in lean mass in mice fed either WD or WD+DEFA5. **D)** Blood glucose concentrations in mice fed either WD or WD+DEFA5 during a glucose tolerance test. **E)** Insulin concentrations in mice fed either WD or WD+DEFA5 during a glucose tolerance test. Data are mean with biological replicates are shown as individual data points or in figure. *** P < 0.001, * P < 0.05

**Supplementary Figure 3.**
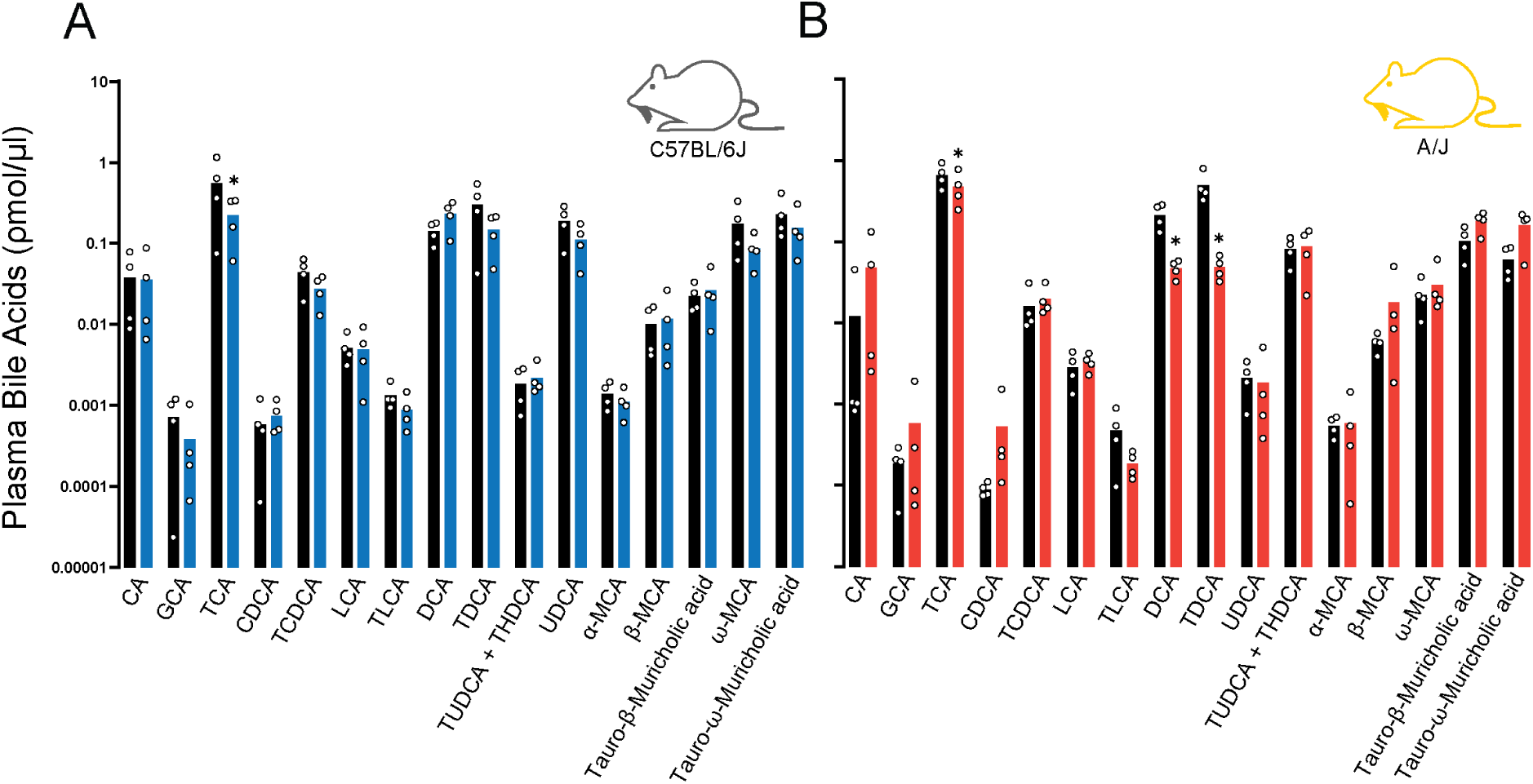
– Effect of alpha-defensin 5 supplementation on insulin sensitivity and body composition in C57BL/6J mice. **A)** Plasma bile acid concentrations from C57BL/6J mice fed either a WD or WD + Defa26 for eight weeks. **B)** Plasma bile acid concentrations from A/J mice fed either a WD or WD + Defa26 for eight weeks. Data are mean with biological replicates are shown as individual data points or in figure. * P < 0.05 denotes a significant difference from WD fed mice.

